# TCF21 loss identifies a conserved inflammation-associated fibroblast state across Crohn’s disease and ulcerative colitis

**DOI:** 10.64898/2026.10.21.760764

**Authors:** Mor Hindi Malowany, Noah Franklin, Matthew V. Russo, Ludovica Arpinati, Yaniv Stein, Adi Grundman, Ruth Scherz-Shouval, Richard L. Gieseck

## Abstract

**Background and Aims:** Inflammatory bowel diseases (IBD), including Crohn’s disease (CD) and ulcerative colitis (UC), are associated with stromal remodeling; however, the fibroblast states that contribute to chronic inflammation remain incompletely defined. We aimed to identify conserved pathogenic fibroblast programs across CD and UC and determine whether these states are linked to loss of homeostatic stromal regulators.

**Methods:** We integrated publicly available single-cell RNA sequencing datasets from healthy and inflamed intestinal tissues to define stromal populations across CD, UC, and control samples. Fibroblast subsets were annotated using cluster-specific markers and patient-level abundance analyses. Multiplex immunofluorescence of human gastrointestinal tissue microarrays assessed protein expression. Human colonic fibroblasts were stimulated with IL1B and TNF to model inflammation-associated fibroblast polarization, and restoration of homeostatic factors was evaluated using mRNA-loaded lipid nanoparticles (LNPs).

**Results:** A conserved inflammation-associated fibroblast (IAF) population was expanded in both CD and UC and expressed inflammatory chemokines, matrix-remodeling genes, and activation markers including *CXCL1*, *MMP3*, *FAP*, and *PDPN*. In contrast, a homeostatic *ADAMDEC1*/*TCF21*-enriched fibroblast population was reduced in inflamed tissue. *TCF21* expression positively correlated with *ADAMDEC1* and negatively correlated with activation markers. Multiplex immunofluorescence confirmed depletion of TCF21-positive non-immune, stromal cells in IBD tissue. *In vitro*, IL1B and TNF induced an IAF-like phenotype and reduced *TCF21*. TCF21 mRNA-LNP restoration attenuated inflammatory mediators including *IL1B* and *IL6*.

**Conclusions:** TCF21 loss identifies a conserved inflammation-associated fibroblast state across CD and UC. Restoring TCF21 partially suppresses inflammatory fibroblast programs, supporting stromal reprogramming as a therapeutic concept in IBD.

## 1. Introduction

Inflammatory bowel diseases (IBD), comprising Crohn’s disease (CD) and ulcerative colitis (UC), are chronic relapsing disorders characterized by dysregulated immune responses, epithelial barrier dysfunction, and progressive tissue remodeling. Despite major advances in biologic therapies, a substantial proportion of patients fail to achieve durable remission, and many develop complications associated with ongoing tissue damage, including fibrosis, stricturing disease, and impaired mucosal healing^1–3^. Although CD and UC differ in their anatomical distribution, depth of tissue involvement, and inflammatory architecture, both ultimately converge on chronic inflammation and disruption of intestinal homeostasis^1–3^. The mechanisms that determine disease progression and therapeutic response remain incompletely understood.

While IBD has historically been viewed through the lens of immune dysregulation, increasing evidence indicates that stromal cells are active participants in disease pathogenesis rather than passive responders to inflammation^4–9^. Stromal cells comprise a complex network of fibroblasts, myofibroblasts, pericytes, smooth muscle cells, and other mesenchymal populations that collectively maintain epithelial integrity, regulate extracellular matrix (ECM) turnover, and coordinate tissue repair. Among these populations, fibroblasts have emerged as key regulators of tissue homeostasis and remodeling^9^. In addition to their established roles in ECM deposition and fibrosis, fibroblasts actively shape local immune responses through the production of cytokines, chemokines, and growth factors that influence both innate and adaptive immunity^9^. Dysregulated fibroblast activation has been implicated in several pathological features of IBD, including fibrosis, stricturing disease, fistula formation, and persistent inflammation^10–12^. Recent single-cell RNA sequencing (scRNA-seq) studies have transformed our understanding of stromal biology in IBD by revealing extensive heterogeneity within intestinal fibroblast populations^5,6,8,13^. Multiple groups have identified stromal subsets enriched in inflamed tissue that express inflammatory chemokines, extracellular matrix-remodeling enzymes, and markers of fibroblast activation, including FAP and PDPN^14–19^. These populations share features with inflammatory and activated fibroblast subsets described in cancer and other chronic inflammatory diseases, suggesting the existence of conserved programs of pathological stromal activation^20–27^. Notably, inflammatory fibroblast populations have been associated with disease severity and resistance to anti-TNF therapy, highlighting their potential importance in driving ongoing tissue pathology^6,7,28–30^. However, the transcriptional mechanisms that maintain fibroblast homeostasis and restrain the acquisition of inflammatory fibroblast phenotypes remain poorly defined. Likewise, whether these inflammatory fibroblast states represent a conserved pathogenic program shared across both CD and UC remains unclear.

While existing therapies primarily target inflammatory cytokines and immune cell function, strategies aimed at restoring stromal homeostasis have received comparatively little attention^9,31–34^. Identifying the molecular regulators that govern fibroblast state transitions may reveal therapeutic opportunities distinct from current immune-directed approaches. Here, we sought to define the stromal landscape of CD and UC and identify conserved fibroblast programs associated with intestinal inflammation. By integrating publicly available scRNA-seq datasets, we identify a conserved inflammation-associated fibroblast (IAF) population that is expanded across both CD and UC. In parallel, we define a homeostatic *ADAMDEC1*/*TCF21*-enriched fibroblast population that is depleted in inflamed tissue. Using human fibroblast models of inflammatory activation, we demonstrate that *TCF21* expression is suppressed during acquisition of an IAF-like phenotype and that restoration of TCF21 partially attenuates key inflammatory programs. Together, these findings identify TCF21 as a candidate regulator of intestinal fibroblast homeostasis and suggest that therapeutic restoration of stromal regulatory pathways may represent a novel approach for the treatment of IBD.

## 2. MATERIALS AND METHODS

### 2.1. Cell lines

The CCD18-Co human colon-derived fibroblast cell line was purchased from ATCC. CCD18-Co cells were expanded and cryopreserved according to ATCC guidelines. Cells were cultured in Dulbecco’s Modified Eagle’s Medium (DMEM) with GlutaMax (Gibco, 10569-010) supplemented with heat-inactivated fetal bovine serum (Gibco, 10082-147) and penicillin-streptomycin (Gibco, 10378016).

### 2.2. Human patient-derived samples

A commercial human colon-disease spectrum tissue microarray (TMA) was purchased from US Biomax Inc. (TissueArray.com; BC05002b). All tissues were obtained by the supplier under HIPAA-approved protocols and with fully informed donor consent. The colon disease spectrum TMA contains 80 cores from 79 cases (1 core/patient plus one core from adrenal gland sample as tissue marker). Each core has a diameter of 1.5 mm and 5 µm thickness. Prior to processing and staining, all FFPE TMA sections were baked for 60 minutes at 60C to prevent tissue detachment.

### 2.3. Multiplexed immunofluorescent (MxIF) staining

FFPE sections from the colon-disease spectrum TMA were de-paraffinized in xylene and fixed in 10% neutral buffered formalin (37% formaldehyde solution diluted in PBS 1x). Antigen retrieval was performed with citrate buffer pH 6.0 (CK18, pan-CK, CK7, EpCAM) or with Tris-EDTA buffer pH 9.0 (PDPN, CD45, TCF21). Following antigen retrieval, FFPE slides underwent photobleaching to minimize background fluorescence, (4.5% w/v H2O2 and 20mM NaOH in PBS) and illuminated using LED lamps. Slides were then blocked using a 10% BSA in PBS solution (BlockerTM BSA, Thermo Scientific). All primary antibodies were diluted in a 2% BSA in 0.05% PBS-T solution (0.05% Tween-20 in PBS) and stained for MxIF imaging using Opal reagents (Akoya Biosciences). Primary antibody incubation overnight at 4°C was followed by incubation with the complementary horseradish peroxidase (HRP) conjugated secondary antibody. Next, TMA slides were incubated with Opal reagents and subsequently washed to perform antigen retrieval as previously explained. Then, slides were incubated with the following primary antibody or with DAPI at the end of the MxIF staining cycle. TMA slides were mounted using Immu-Mount (Epredia, #9990402). All antibodies were first validated and optimized individually. The following primary antibody sequence was used for the final TMA MxIF staining as shown in Figure 2: Epithelial cell cocktail (EpCAM + Cytokeratin 18 + Cytokeratin 7 + pan-Cytokeratin) → PDPN → CD45 → TCF21 → DAPI. Slides were imaged using PhenoImager HT scanner (Akoya Biosciences) and analyzed using QuPath (version 0.5.1). Detailed antibody information: Anti-Cytokeratin 18 (Abcam; Cat# ab7797; clone DC10), anti-Cytokeratin 7 (Abcam; Cat# ab181598; clone EPR17078), anti-pan-Cytokeratin (Dako; Cat# M3515; clone AE1/AE3), anti-CD326/EpCAM (Cell Signaling Technology; Cat# 14452; D9S3P), anti-PDPN (Abcam; Cat# ab236529; clone EPR22182), anti-CD45 (Cell Signaling Technology; Cat# 13917; D9M8I), anti-TCF21 (Creative Diagnostics; Cat# DPABH-15598; aa42-91).

### 2.4. MxIF analysis

For image background removal, the Akoya spectral unmixing algorithm was used during image post processing based on a non-stained slide. Images were analyzed using QuPath version 0.5.1, and ImageJ-Fiji for snapshots and single-channel representative images. TMA cores were defined using the QuPath TMA de-array tool. Cores displaying less than one third of tissue due to either damage in processing or staining were excluded from the analysis. Upon relevant TMA core selection, an object-based analysis was performed. Cell segmentation was carried out using the integrated “cell detection” tool in QuPath version 0.5.1, based on nuclear staining (DAPI). Subsequently, we employed a machine learning approach for marker classification. The “Random Trees” classifier was trained to define cells as positive for the different markers, or to ignore them. This object-based marker classification was performed independently for each fluorescent channel of interest, in addition to the DAPI channel. Once each marker classifier was trained, we applied a composite classifier comprising all marker classifiers to the whole TMA image. Fluorescent intensity measures and cell count data were extrapolated for all possible marker combinations. Positive cells for a selected marker represent the percentage of signal-positive cells per core, relative to the total cell count in each core. Using the object-based classification, we defined epithelial cells as EPCAM^+^CD45^-^PDPN^-^, immune cells as EPCAM^-^CD45^+^PDPN^-^, PDPN^+^ fibroblasts as EPCAM^-^CD45^-^PDPN^+^, and PDPN^-^stroma as EPCAM^-^CD45^-^PDPN^-^. For more general quantifications, we defined epithelial cells as EPCAM^+^CD45^-^, immune cells as EPCAM^-^CD45^+^ and non-immune stroma as EPCAM^-^CD45^-^.

### 2.5. Induction of inflammatory-like phenotype *in vitro*

CCD-18Co cells were plated in a 96-well format at a density of 20,000 cells/well and incubated overnight to adhere. The following day, media containing 1ng/ml IL1B (R&D Systems, 201-LB-005/CF), 1ng/ml TNF (R&D Systems, 210-TA-020/CF), or both was added to designated wells and the cells were stimulated for an additional 4hrs (gene expression) or 24hrs (protein detection). After stimulation, supernatants were collected and frozen at -80C until analyzed and cells were lysed in RLT buffer (Qiagen) and stored at -80C until processed for RNA.

### 2.6. mRNA-loaded lipid nanoparticles (LNP) for gene modulation

CCD-18Co cells were seeded onto 24-well plates at a concentration of 0.35×10^5^ cells/well and left to adhere overnight. The following day, medium was replaced with DMEM full medium, and cells were cultured in the presence or absence of the inflammatory cytokine cocktail (0.5 ng/mL of TNF, ThermoFisher, 300-01A-50); and 0.05 ng/mL of IL1B, ThermoFisher, 200-01B-50). 3 hours post inflammation induction, LNP-hTCF21-P2A-EGFP or empty vector control EGFP-LNP-mRNA were added to the cells at a working concentration of 1 μg/mL. Following 24 hours treatment, cells were harvested and processed for RT-qPCR analysis or immunofluorescence staining (IF). Custom-made lipid nanoparticles (LNP) containing human *TCF21* mRNA (LNP-hTCF21-P2A-EGFP mRNA, m1Psi modified) for *in vitro* transcription (IVT), and empty vector controls (EGFP-LNP-mRNA, cat# NL1011-0100) were purchased from VectorBuilder Inc., Chicago, USA.). hTCF21-P2A-EGFP sequence of interest and LNP details: ATGTCCACCGGCTCCCTCAGCGATGTGGAGGACCTTCAAGAGGTGGAGATGTTGG AATGTGACGGGTTGAAAATGGATTCGAACAAGGAATTTGTGACTTCCAACGAGAGCACCGAGGAGAGCTCCAACTGCGAGAATGGGTCTCCCCAGAAGGGCCGCGGCGGC CTGGGCAAGAGGAGGAAGGCGCCCACCAAGAAGAGCCCCCTGAGCGGGGTCAGCCAGGAGGGGAAGCAGGTCCAGCGCAACGCCGCCAACGCGCGAGAGCGGGCCCGCATGCGAGTGCTGAGCAAGGCCTTCTCCAGACTCAAGACCACCCTGCCCTGGGTGCCCCCCGACACCAAGCTCTCCAAGCTGGACACGCTCAGGCTGGCGTCCAGCTACATCGCCCACTTGAGGCAGATCCTGGCTAACGACAAATACGAGAACGGGTACATTCACCCGGTCAACCTGACGTGGCCCTTTATGGTGGCCGGGAAACCCGAGAGTGACCTGAAAGAAGTGGTGACCGCGAGCCGCTTATGTGGAACCACCGCGTCCGGAAGCGGAGCCACGAACTTCTCTCTGTTAAAGCAAGCAGGAGATGTTGAAGAAAACCCCGGGCCTATGGTGAGCAAGGGCGAGGAGCTGTTCACCGGGGTGGTGCCCATCCTGGTCGAGCTGGACGGCGACGTAAACGGCCACAAGTTCAGCGTGTCCGGCGAGGGCGAGGGCGATGCCACCTACGGCAAGCTGACCCTGAAGTTCATCTGCACCACCGGCAAGCTGCCCGTGCCCTGGCCCACCCTCGTGACCACCCTGACCTACGGCGTGCAGTGCTTCAGCCGCTACCCCGACCACATGAAGCAGCACGACTTCTTCAAGTCCGCCATGCCCGAAGGCTACGTCCAGGAGCGCACCATCTTCTTCAAGGACGACGGCAACTACAAGACCCGCGCCGAGGTGAAGTTCGAGGGCGACACCCTGGTGAACCGCATCGAGCTGAAGGGCATCGACTTCAAGGAGGACGGCAACATCCTGGGGCACAAGCTGGAGTACAACTACAACAGCCACAACGTCTATATCATGGCCGACAAGCAGAAGAACGGCATCAAGGTGAACTTCAAGATCCGCCACAACATCGAGGACGGCAGCGTGCAGCTCGCCGACCACTACCAGCAGAACACCCCCATCGGCGACGGCCCCGTGCTGCTGCCCGACAACCACTACCTGAGCACCCAGTCCGCCCTGAGCAAAGACCCCAACGAGAAGCGCGATCACATGGTCCTGCTGGAGTTCGTGACCGCCGCCGGGATCACTCTCGGCATGGACGAGCTGTACAAGTAA.

Technical details of the LNPs: Polidispersity index (PDI) = 0.1, Zeta potential = -2.7 mV, particle size = 94.37 nm.

### 2.7. In vitro Immunofluorescent (IF) staining

CCD-18Co cells expressing LNP-hTCF21-P2A-EGFP or empty vector control EGFP-LNP-mRNA were fixed with 2% PFA solution for 30 minutes at room temperature and permeabilized with 0.5% Triton-X 100 in blocking buffer (PBS containing 5% BSA) for 1 hour at room temperature. Then, CCD-18Co cells were stained with primary TCF21 antibody (Creative Diagnostics; Cat# DPABH-15598; aa42-91) overnight at 4°C. The next day, cells were incubated with secondary AlexaFluor-568 antibody for 1 hour at room temperature and subsequently stained for DAPI before being mounted. Images were acquired using Nikon Eclipse Ci microscope and were analyzed using ImageJ-Fiji software.

### 2.8. Cell identification

Cluster-specific marker genes for stromal subpopulations were identified using the U-method package . Uniquely expressed genes (UEGs) were defined based on expression probability differences across clusters, and statistical significance was assessed using Benjamini-Hochberg-adjusted p-values, with a threshold of adjusted p-value < 0.05.

### 2.9. qRT-PCR

After 4hrs of cytokine stimulation, supernatant was aspirated and the cells were washed 2X with PBS. After aspirating off the PBS, the cells were directly lysed in the wells by the addition of 350ul RLT buffer (Qiagen). Purified RNA was collected by using an RNAeasy Kit (Qiagen) and quantified using the BioTek Synergy instrument. Purified RNA (∼20ng) was used to detect gene expression with the TaqMan RNA-to-Ct 1-Step Kit (Life Technologies). Briefly, reverse transcription and real-time quantitative PCR were performed in the same well on a QuantStudio 7 instrument using this kit. Gene expression for *CXCL1* (Taqman, Hs00236937_m1) and *CXCL8* (Taqman, Hs00174103_m1) were calculated using normalization to *RPLP0* as the housekeeping gene. Fold changes over untreated samples were determined by calculating the delta CT and delta-delta CT values based on the *RPLP0* expression per sample.

### 2.10. Secreted protein detection by Luminex

After 24hrs of cytokine stimulation, supernatant samples were collected by pipetting and stored at -80C until the day of analysis. We used the Human Immune Monitoring Procarta Luminex panel (ThermoFisher) to detect 65 analytes in the supernatant samples. Briefly, each sample was spun down for 1 min @ 1,000g to remove any remaining particulates. Every sample was diluted 1:5 in cell media and incubated with the analyte bead mixture for 2hrs at room temperature on an orbital shaker set to 500rpm. After washing via magnetic plate, detection antibodies were added and samples were incubated for 30mins on the orbital shaker. After a second washing, streptavidin-PE bead mixture was added and the samples were incubated for a final 30mins on the orbital shaker. After a final wash, the samples were analyzed on the FlexMap 3D instrument. FlexMap software was used to calculate standard curves for each of the 65 analytes and quantify expression of each for every sample. Due to the large number of analytes and samples, an R programming script (see Supplementary File) was used to import the raw data and format it into heatmap visualizations.

### 2.11. Single cell RNA sequencing

Fixed cells were rehydrated as described in the 10x Genomics methanol fixation protocol (CG000136 Rev E) and labelled with custom cholesterol-modified oligonucleotides of the MULTI-seq design^35^. Each CCD-18Co replicate received a unique barcode. Labelled samples were combined into a mixed CCD-18Co pool. The CCD-18Co pool was loaded into a single Chromium reaction targeting 15,000 cells. Gene expression libraries were prepared according to the Chromium Next GEM Single Cell 3′ Gene Expression v3.1 (Dual Index) protocol (CG000315 Rev C) with Dual Index Kit TT Set A. Barcode libraries were recovered from the cDNA cleanup supernatant and amplified for 6 cycles using TruSeq small RNA (RPI) indexing primers. Libraries were pooled at approximately 96:4 (gene expression:barcode) and sequenced on an Illumina NextSeq 500 High Output flow cell (28/10/10/91), targeting 5,000 and 1,250 reads per cell, respectively. FASTQs were generated using cellranger mkfastq (v6.0.0, 10x Genomics) on Seven Bridges, run separately for each library type.

### 2.12. Statistical analysis

Statistical analysis and visualization were conducted utilizing GraphPad Prism 10 (GraphPad; RRID: SCR_002798). To analyze normally distributed data, Welch’s t-test was used for comparisons between two groups, and one-way ANOVA with appropriate post-hoc testing for multiple comparisons. *p* value < 0.05 or = FDR <0.05 were used as parameters to determine statistical significance for *t* test and multiple comparisons, respectively. * = *p*<0.05; **= *p*<0.01; *** = *p*<0.001. Non-significant differences (*p* value ≥ 0.05) are indicated with “ns”.

### 2.12. IBD Stromal Atlas Construction and Analysis

Seurat package^36^ was used for processing, integrating, normalizing, and analyzing data from the studies referenced here^6,8,13,37^. Cells were filtered to ensure > 200 genes per cell and mitochondrial counts < 30%. Ribosomal genes were removed and any cells expressing *PTPRC*, *IGLL1*, or *EPCAM* were removed to enrich for stromal fractions. Counts were normalized using SCTransform, regressing on mitochondrial percentage. Harmony package^38^ was used to correct for variations in chemistry used for library generation and source of data. Clustree package was used to assess clustering from a resolution of 0.05 to 1.0 using the first 50 principal components. A resolution of 0.1 was ultimately used to define cell types.

Regional identity of samples was integrated into a harmonized framework as follows: “ILEUM”, “Ileum”, “Ileal”, “SmallInt” and "TI" were annotated as “Small Intestine”. "Colon",

“Right_Colon”, “Left_Colon”, “Transverse_Colon”, “Sigmoid_Colon”, “Ascending”, “Desc ending”, “Transverse”, “Sigmoid”, “Caecum”, “Cecum”, “Proximal_Sigmoid”, “Rectum”, “APD”, “REC”, and “LargeInt” were annotated as “Colon”. “lymph node” and “#N/A” were annotated as “Unknown” and dropped from subsequent analyses.

Disease annotation of samples was integrated into a harmonized framework as follows: “Uninvolved.iCD”, “Involved.iCD”, “CD”, “Pediatric Crohn Disease”, “CD.Non-inflamed”, “CD.Non-stricture”, and “CD.Stricture” were annotated as “Crohn’s Disease”. “Inflamed”, “Non-inflamed”, “UC”, “UC.UC.Inflamed”, and “UC.UC.Non-inflamed” were annotated as “Ulcerative Colitis”. “Healthy”, “Control”, “Pediatric healthy”, “Healthy adult”, and “Normal” were annotated as “Healthy”.

Top differential marker genes among clusters were identified using the package UMethod^39^ with the parameters Uscore = 0.2 and n = 5. These were used to assign identities to the cell clusters, after which contaminating epithelial cells and dying / apoptotic cells were removed.

For patient level analyses, patients with <250 total cells were removed to prevent bias in data. Unpaired T tests were used to calculate statistical significance for differences in cell type abundance between disease conditions.

### 2.13. Use of AI writing assistance

During the preparation of this work, the authors used Microsoft Copilot to improve the readability and language of the manuscript and assist in formatting. After using this tool/service, the authors reviewed and edited the content as needed and take full responsibility for the content of the published article.

## 3. Results

### 3.1. A conserved inflammation-associated fibroblast population is expanded across both Crohn’s disease and ulcerative colitis

To determine how stromal populations are altered during intestinal inflammation, we integrated stromal cells from four independent scRNA-seq datasets^6,8,13,37^ encompassing healthy intestine, ulcerative colitis (UC), and Crohn’s disease (CD). After quality control and filtering, 104,581 stromal cells derived from 89 unique samples were included for analysis, with 11.0% originating from healthy or normal-adjacent colon, 15.4% from healthy or normal-adjacent small intestine, 14.5% from lesional UC colon, and 59.1% from lesional CD small intestine (Figure 1A). To account for variation in sample processing, sequencing chemistry, and study origin, datasets were integrated using *Harmony*^38^ batch correction (Figure 1B). Low-resolution clustering identified twelve stromal populations, which were further refined following removal of contaminating epithelial cells, B cells, and apoptotic cells, yielding nine major stromal populations for downstream analyses (Figures 1C-D, S1A-B).

**Figure 1.**
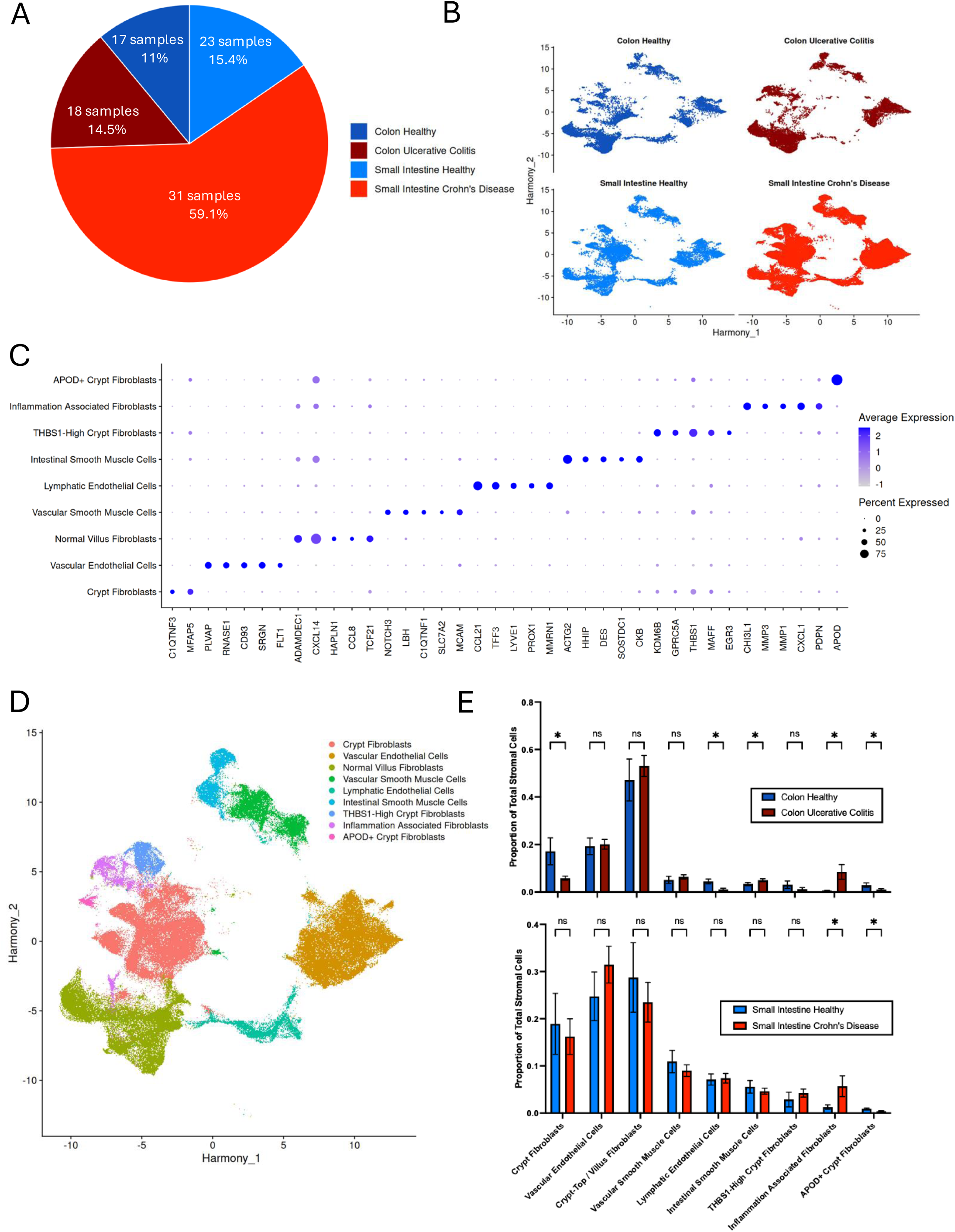
A conserved inflammation-associated fibroblast population is expanded across Crohn’s disease and ulcerative colitis. (A) Composition of the integrated stromal atlas showing the distribution of sample counts and total cell percentages derived from healthy intestine, colon, ulcerative colitis (UC), and Crohn’s disease (CD) samples. (B) Harmony-corrected UMAP visualization of integrated stromal cells derived from four independent scRNA-seq datasets. (C) Expression dotplot showing expression of representative marker genes used for assignment of stromal identities. (D) UMAP projection of major stromal populations identified following clustering and annotation. (E) Quantification of stromal population abundance across disease states. Inflammation-associated fibroblasts (IAFs) are significantly expanded in both UC and CD compared with healthy controls, whereas crypt fibroblasts and lymphatic endothelial cells are reduced. Statistics are representative of one-sided unpaired Welch’s t-test with significance shown at p < 0.05.

To annotate these populations, we applied the *U-method*^40^ to identify uniquely expressed genes within each cluster. Consistent with previous descriptions of intestinal stromal heterogeneity, we identified endothelial, lymphatic endothelial, smooth muscle, and fibroblast populations (Figure 1C). Among fibroblasts, one population expressed high levels of inflammatory chemokines, matrix-remodeling enzymes, and fibroblast activation markers, including *CXCL1*, *MMP1*, *MMP3*, *FAP*, and *PDPN*, consistent with previously described inflammation-associated fibroblasts (IAFs)^9^ (Figure 1C-D).

We next quantified the relative abundance of each stromal population across disease states. Crypt fibroblasts and lymphatic endothelial cells were significantly depleted in inflamed colon tissues, whereas IAFs were markedly expanded in both UC and CD (Figure 1E). In healthy tissues, IAFs comprised approximately 1% of stromal cells, whereas they expanded to more than 5% of stromal cells in CD and 10% in UC. These findings demonstrate that a transcriptionally conserved inflammation-associated fibroblast population emerges across both major forms of IBD despite their distinct clinical and anatomical manifestations.

### 3.2. TCF21 is anti-correlated with IAF phenotype

Prior studies have implicated ADAMDEC1 in protection from intestinal inflammation and maintenance of mucosal homeostasis^41^. Within the *ADAMDEC1*-enriched fibroblast population, *TCF21* emerged as the top enriched transcription factor, suggesting a potential role in maintenance of fibroblast identity. TCF21 is a basic helix-loop-helix transcription factor implicated in mesenchymal cell differentiation and fibroblast quiescence in several tissues, although its role within the intestinal stroma remains poorly understood^42–47^. Compared with other intestinal fibroblast populations, IAFs expressed markedly lower levels of *ADAMDEC1*, irrespective of disease status (Figure 2A). Notably, *TCF21* expression closely tracked with *ADAMDEC1* expression, displaying significant positive correlations in healthy intestine, UC, and CD samples (Figure 2B).

**Figure 2.**
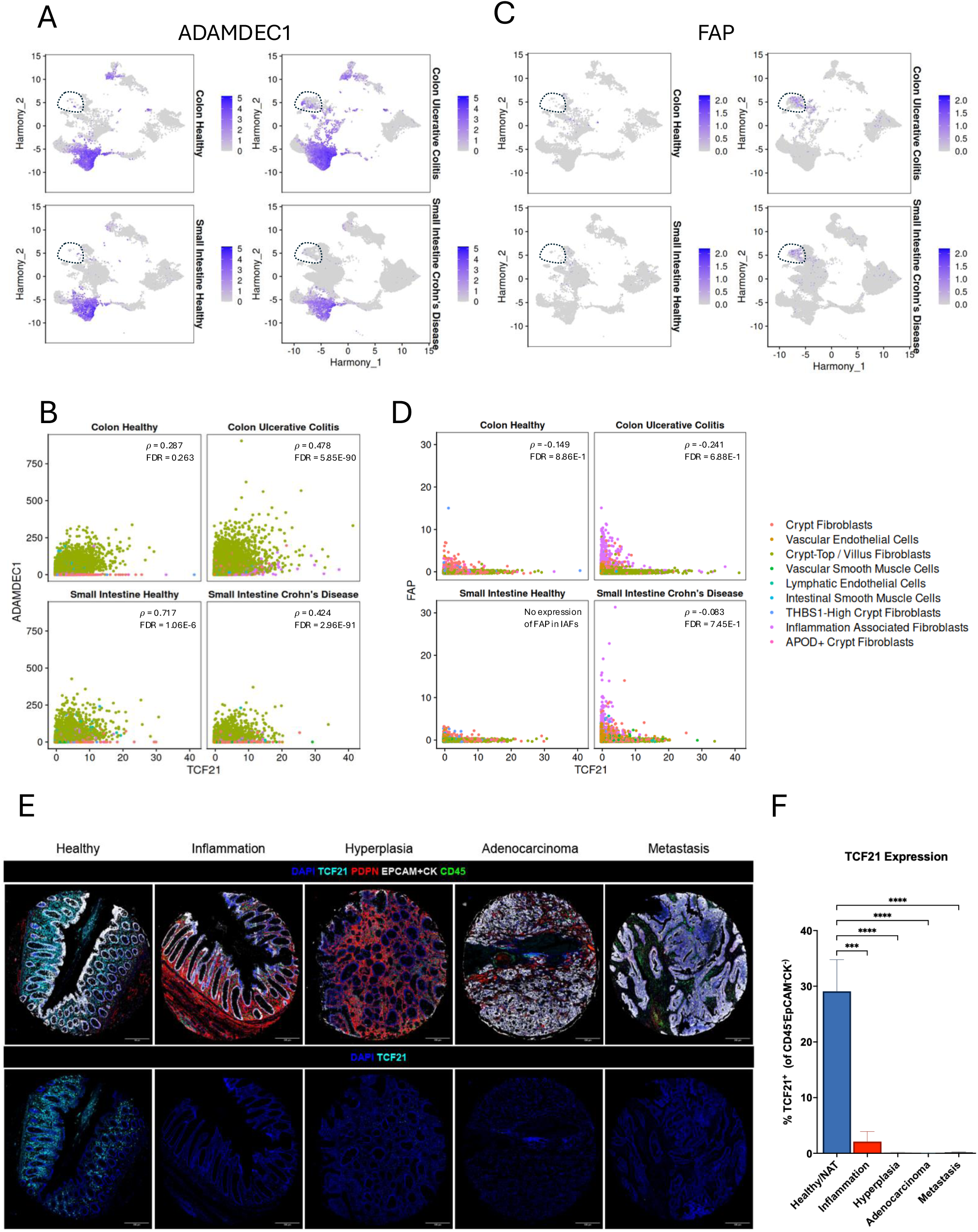
TCF21 marks a homeostatic fibroblast population that is depleted during intestinal inflammation. (A) Harmony corrected UMAPs showing expression of *ADAMDEC1* across stromal populations. (B) Correlation between *ADAMDEC1* and *TCF21* expression in healthy colon, intestine, UC, and CD samples. (C) Harmony corrected UMAPs showing expression of *FAP* across stromal populations. (D) Correlation between *FAP* and *TCF21* expression in healthy colon, intestine, UC, and CD samples. (E) Representative multiplex immunofluorescence images of healthy, inflammatory bowel disease, tumor-bearing, and metastatic colorectal cancer tissues stained for epithelial markers, CD45, PDPN, and TCF21. (F) Quantification of TCF21-positive non-immune stromal cells across tissue groups. A total of 63 cores were included in the analysis: Healthy/NAT (n=16), Inflammation (n=6), Hyperplasia (n=6), Adenocarcinoma (n=17), and Metastasis (n=18). Statistical analysis was performed using one-way ANOVA with adequate post-hoc testing. Bar plots shown as mean±SD *** p<0.05, ****p<0.01.

In contrast, expression of the canonical activation marker *FAP* exhibited the opposite relationship. IAFs were enriched for *FAP*, particularly within inflamed tissues (Figure 2C), and *FAP* levels were significantly negatively correlated with *TCF21* expression across healthy and diseased samples (Figure 2D).

To determine whether these transcriptional findings were reflected at the protein level, we performed multiplex immunofluorescence analysis on gastrointestinal tissue microarrays containing healthy, inflamed, tumor-bearing, and metastatic colorectal cancer specimens (Figure 2E). In healthy and normal-adjacent tissues, approximately 30% of non-immune stromal cells expressed TCF21 (Figure 2F). In contrast, TCF21-positive stromal cells were markedly depleted in IBD tissues, with only 2% of stromal cells retaining expression. Because inflammation-associated fibroblasts share several features with cancer-associated fibroblasts, we next examined whether TCF21 loss extended beyond chronic inflammation. Consistent with this possibility, TCF21 expression was further reduced in tumor-associated stroma and metastatic colorectal cancer tissues (Figures 2E-F). Collectively, these findings identify TCF21 as a marker of a homeostatic fibroblast population that is progressively lost during intestinal inflammation and neoplastic transformation.

### 3.3. Inflammatory cytokines induce an IAF-like phenotype and suppress *TCF21* expression *in vitro*

The reciprocal relationship between TCF21 and inflammatory fibroblast markers suggested that inflammatory signaling may actively drive loss of TCF21 during fibroblast activation. To test this hypothesis, we established an *in vitro* model of inflammatory fibroblast polarization using CCD-18Co human colonic fibroblasts stimulated with IL1B and TNF (Figure 3A). At four hours following stimulation, both IL1B and TNF induced robust upregulation of the inflammatory chemokines *CXCL1* and *CXCL8*, with IL1B producing the strongest effects (Figure 3B). Combined IL1B/TNF stimulation did not further augment chemokine expression, instead generating responses comparable to IL1B alone. Analysis of secreted proteins after 24 hours using multiplex Luminex profiling demonstrated distinct cytokine-specific responses, with the combined condition again closely resembling the IL1B-driven profile (Figure 3C).

**Figure 3.**
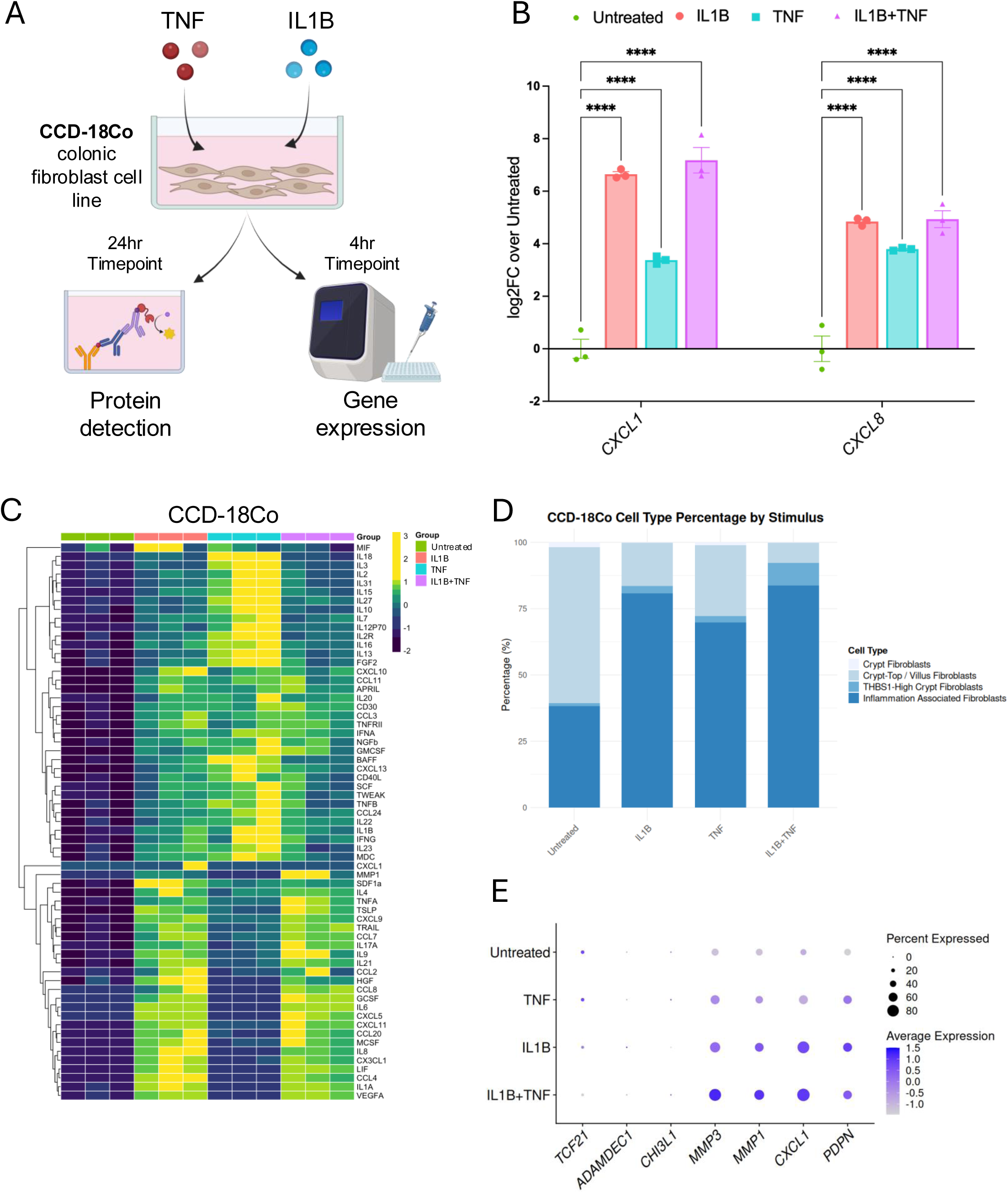
Inflammatory cytokines induce an inflammation-associated fibroblast phenotype in vitro. (A) Experimental design for inflammatory fibroblast polarization using CCD-18Co cells stimulated with IL1B, TNF, or the combination of both cytokines. (B) qPCR analysis of *CXCL1* and *CXCL8* following cytokine stimulation. (C) Heatmap summarizing secreted cytokines and chemokines measured by multiplex Luminex analysis. (D) Projection of stimulated fibroblasts onto the human stromal atlas (Figure 1). Cytokine stimulation results in a shift from homeostatic fibroblast populations toward the inflammation-associated fibroblast compartment. (E) Dotplot expression of hallmark IAF-associated genes and *TCF21* following cytokine stimulation.

To determine how closely this model recapitulates fibroblast states observed in human disease, we performed single-cell RNA sequencing on stimulated fibroblasts and projected these profiles onto our human stromal atlas. Under basal conditions, approximately 60% of CCD-18Co cells mapped to the normal villus fibroblast compartment, whereas the remaining cells displayed partial similarity to inflammatory fibroblasts (Figure 3D). Following cytokine stimulation, however, the majority of cells mapped to the IAF compartment. IL1B alone and IL1B/TNF combination treatment produced the strongest shift toward this inflammatory state, demonstrating that inflammatory cytokine exposure recapitulates key transcriptional features of patient-derived IAFs.

Finally, we examined expression of hallmark IAF genes following cytokine stimulation. With the exception of *CHI3L1*, which was minimally expressed in CCD-18Co cells, canonical IAF markers including *MMP1, MMP3, CXCL1,* and *PDPN* were significantly induced following stimulation (Figure 3E). Additionally, acquisition of this inflammatory phenotype was accompanied by a concomitant reduction in *TCF21* expression. Together, these findings demonstrate that inflammatory cytokines are sufficient to drive an IAF-like transcriptional program while simultaneously suppressing expression of the homeostatic regulator TCF21.

### 3.4. Restoration of TCF21 attenuates inflammatory fibroblast programs

The depletion of TCF21 in patient samples and during cytokine-induced fibroblast activation suggested that loss of TCF21 may contribute to maintenance of inflammatory fibroblast states. We therefore hypothesized that restoring TCF21 expression would attenuate inflammatory fibroblast programs and partially restore features of fibroblast homeostasis. To calibrate our experimental system and ensure a physiologically relevant setting for targeting TCF21, we performed a dose-response analysis on CCD-18Co cells using TNF and IL1B stimulation. This analysis demonstrated a concentration-dependent upregulation of established inflammatory markers (CXCL1, IL1β) alongside a progressive suppression of endogenous TCF21 (Figure 4A, B). To assess whether restoring TCF21 expression in inflammatory fibroblasts could induce a shift towards their original homeostatic phenotype, we generated lipid nanoparticles (LNPs) encapsulating TCF21 mRNA and administered these particles to fibroblasts undergoing inflammatory activation (Figures 4B-E). Treatment with TCF21-LNP resulted in robust expression of *TCF21* (Figure 4C), increasing transcript levels by approximately 100-fold relative to controls (Figure 4D). Restoration of TCF21 was accompanied by significant reductions in expression of the inflammatory mediators *IL1B* and *IL6* (Figures 4E-F), demonstrating that manipulation of fibroblast identity can influence downstream inflammatory programs.

**Figure 4.**
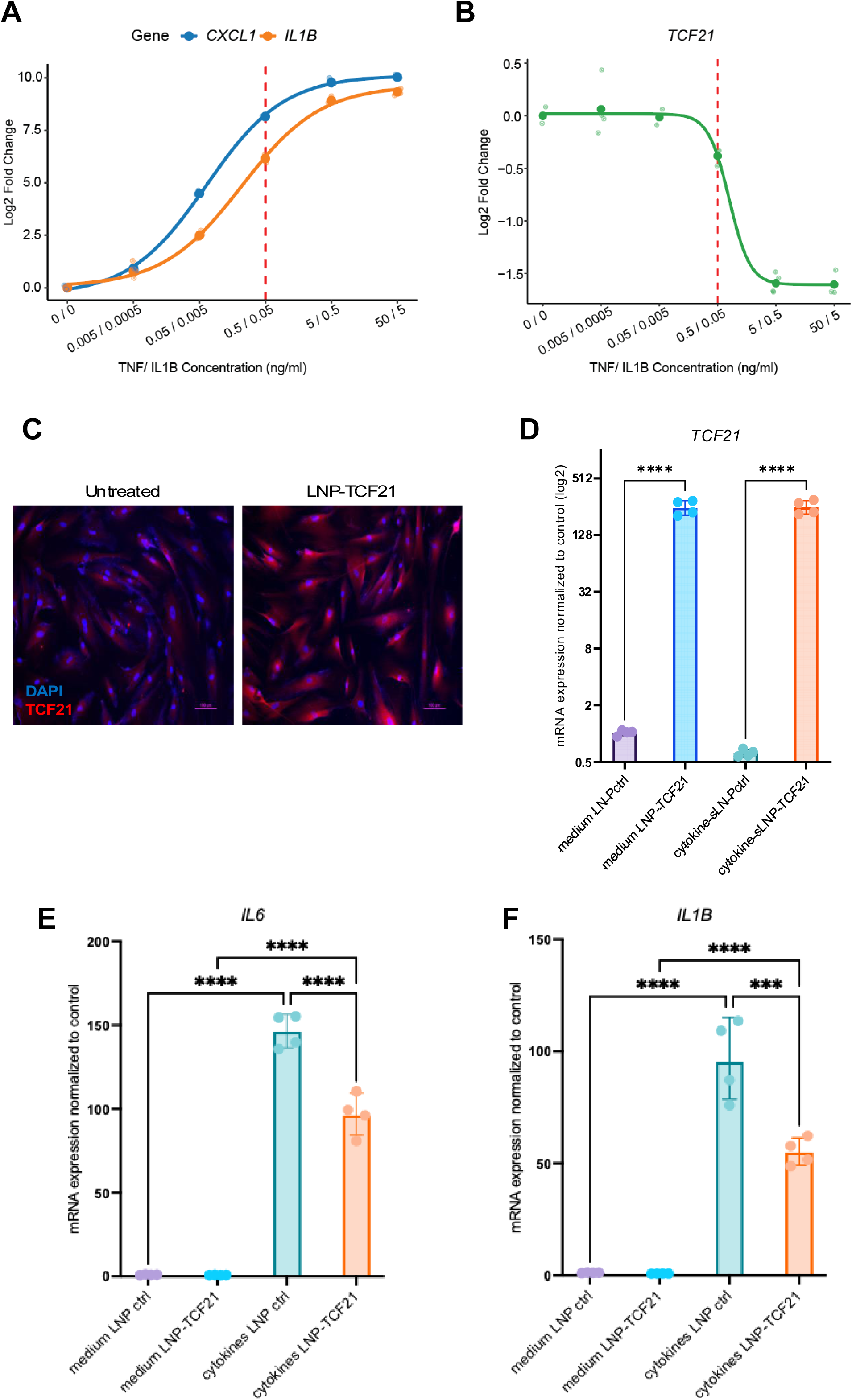
TCF21 restoration suppresses inflammatory signaling in activated fibroblasts. (A-B) Log2 fold expression changes of the inflammatory mediators CXCL1 and IL1B (A), and *TCF21* (B), in CCD-18Co cells, following a 48 hr dose-response stimulation with TNF and IL1B. The vertical red dashed line indicates the chosen working concentration (0.5/0.05 ng/ml). (C) Immunofluorescence staining validating the upregulation of TCF21 protein levels following 24 hr TCF21-LNP transfection in CCD-18co cells. In blue: DAPI nuclear staining, in red: TCF21 protein. qRT-PCR analysis showing *TCF21* overexpression (D), *IL6* (E), and *IL1B* (F) following TCF21-LNP transfection in CCD-18co cells in both steady state (medium+LNP) and inflammation (cytokines + LNPs). Bar plots shown as mean±SD. ***p<0.05, ****p<0.01.

Importantly, TCF21-mediated suppression of inflammatory signaling remained detectable even when LNPs were administered after inflammatory stimulation had already begun. Although the magnitude of this effect was modest, both IL1B and IL6 were significantly reduced under these conditions (Figures 4E-F), suggesting that activated fibroblasts retain at least partial plasticity. Collectively, these findings establish TCF21 as a disease-modulated regulator of intestinal fibroblast state. While restoration of TCF21 did not fully reverse inflammatory activation, it attenuated key inflammatory programs and provides proof-of-concept that fibroblast identity can be pharmacologically manipulated through transcription factor replacement.

## 4. Discussion

Despite major advances in biologic therapies, progressive tissue remodeling remains a major driver of pathology in both Crohn’s disease (CD) and ulcerative colitis (UC). Inflammatory fibroblasts have emerged as important contributors to intestinal inflammation and therapeutic non-response, yet whether these cells represent disease-specific programs or a shared pathogenic state across IBD has remained unclear^48^. Here, we define the stromal landscape of the small and large intestine across health, CD, and UC and identify a transcriptionally conserved inflammation-associated fibroblast (IAF) population enriched in both diseases. In parallel, we identify a homeostatic *ADAMDEC1*^+^/*TCF21*^+^ fibroblast population that is progressively depleted during inflammation and demonstrate that restoration of TCF21 can partially suppress inflammatory fibroblast programs. Together, these findings suggest that loss of fibroblast homeostasis is a central feature of IBD and identify TCF21 as a candidate regulator of this process.

Fibroblasts are increasingly recognized as plastic cells capable of transitioning between homeostatic and activated states in response to environmental cues, a concept well established in cancer through the description of inflammatory and matrix-remodeling cancer-associated fibroblast (CAF) subsets^20,21^. Our findings extend this framework to IBD by demonstrating that the IAF signature, marked by *CXCL1*, *MMP1*, *MMP3*, *FAP*, and *PDPN*, is not restricted to a single disease entity or anatomical location, but instead emerges consistently across ulcerative colitis colon and Crohn’s disease small intestine.

Although ADAMDEC1 has previously been linked to protection from intestinal inflammation and maintenance of mucosal homeostasis^41^, the upstream regulatory mechanisms governing this fibroblast state have remained poorly understood. Our analyses identified TCF21 as the most prominent transcription factor associated with this homeostatic fibroblast population. TCF21 has emerged as an important regulator of mesenchymal cell identity and fibroblast quiescence in several tissues, particularly within the cardiovascular system, where loss of TCF21 promotes transition toward activated myofibroblast states^44^. Our findings suggest that a similar regulatory axis may operate within the intestinal stroma.

Notably, TCF21^+^ fibroblasts declined progressively across the inflammation-to-neoplasia continuum, decreasing from approximately 30% of stromal cells in healthy tissues to less than 2% in inflamed IBD samples. Although these observations do not establish causality, they raise the possibility that emergence of inflammatory fibroblasts reflects loss of a homeostatic fibroblast transcriptional program. In this model, chronic inflammatory signals may drive progressive erosion of fibroblast identity, permitting acquisition of inflammatory and tissue-remodeling functions that contribute to disease persistence. This concept aligns with emerging views of fibroblast plasticity in both inflammatory disease and cancer and suggests that restoration of homeostatic stromal programs could represent a therapeutic strategy distinct from direct suppression of inflammatory cytokine pathways.

To explore this possibility, we utilized an mRNA-loaded lipid nanoparticle (LNP) platform to restore TCF21 expression in inflammatory fibroblasts. Fibroblasts have historically proven challenging to genetically manipulate, creating a significant obstacle for stromal-directed therapeutic approaches^49,50^. Using this platform, we achieved robust TCF21 overexpression in colonic fibroblasts and observed suppression of key inflammatory mediators, including IL1B and IL6. Importantly, these effects remained detectable even when TCF21 was introduced after inflammatory priming, suggesting that pathogenic fibroblast states retain at least some degree of reversibility. These findings provide proof-of-concept that transcriptional regulators governing fibroblast identity can be manipulated pharmacologically and support the broader concept of stromal reprogramming as a therapeutic strategy in chronic inflammatory disease.

Importantly, restoration of TCF21 did not fully revert inflammatory fibroblasts to a homeostatic phenotype. While expression of several inflammatory mediators was reduced, many features of the activated state persisted, indicating that TCF21 restoration alone is unlikely to completely reverse established fibroblast activation. This incomplete reprogramming suggests that inflammatory fibroblast identity is maintained by multiple convergent pathways and may exhibit features of transcriptional or epigenetic memory. Consequently, therapeutic strategies aimed at restoring fibroblast homeostasis may ultimately require combination approaches involving both upstream cytokine blockade and downstream stromal reprogramming. Such approaches could simultaneously reduce inflammatory signals while promoting re-establishment of a healthy mesenchymal compartment capable of supporting tissue repair and mucosal healing.

Several limitations should be considered when interpreting these findings. First, our stromal atlas was generated through integration of publicly available single-cell datasets collected across different patient populations, tissue processing methods, and sequencing platforms. Although the identification of both IAF and *ADAMDEC1*^+^/*TCF21*^+^ populations was highly consistent across datasets, technical and clinical heterogeneity may influence estimates of subtype abundance. Second, mechanistic validation was primarily performed using an immortalized colonic fibroblast cell line rather than primary patient-derived fibroblasts. Third, while TCF21 restoration attenuated inflammatory gene expression, the downstream transcriptional networks directly regulated by TCF21 in intestinal fibroblasts remain incompletely characterized. Finally, the therapeutic potential of TCF21 restoration will require validation in preclinical *in vivo* models capable of assessing effects on inflammation, fibrosis, tissue remodeling, and mucosal healing.

Collectively, these findings support a model in which chronic intestinal inflammation is accompanied by depletion of a homeostatic TCF21^+^ fibroblast compartment and expansion of a conserved inflammation-associated fibroblast state shared across both CD and UC. Beyond identifying TCF21 as a candidate regulator of this transition, our study highlights stromal reprogramming as a complementary therapeutic strategy to existing immune-directed approaches. As increasing attention is focused on durable disease modification, tissue restoration, and mucosal healing in IBD, interventions that normalize fibroblast states may offer a new avenue for limiting progressive tissue damage and improving long-term outcomes for patients.

## Acknowledgements

The authors would like to acknowledge Josh Fienmman, Ying Zhang, and the Pfizer Next-Generation Sequencing Technology Center for assistance in sample and library preparation. Additionally, the authors thank Liat Alyagor and Daniela Blank from the Department of Veterinary Resources at the Weizmann institute of Science, for their technical assistance with MxIF TMA scanning on the PhenoImager, supported by a research grant from the Quinquin Foundation. L. Arpinati was supported by the Lombroso family fund fellowship. R. Scherz-Shouval is incumbent of The Robert and Yadelle Sklare Professorial Chair in Biochemistry.

## Author Contributions

N.F.: was involved in the design of the study, experimentation, analysis of data, and the conception and writing of the manuscript. M.H.M.: was involved in the design of the study, experimentation, analysis of data, conception and writing of the manuscript. M.V.R.: was involved in the design of the study, experimentation, analysis of data, conception and writing of the manuscript. Y.S.: was involved in the design of the study, analysis of data, conception and writing of the manuscript. L.A.: was involved in the design of the study, experimentation, and analysis of data. A.G.: was involved in the design of the study, experimentation, and analysis of data. R.S.: was involved in the design of the study, conception and writing of the manuscript. R.L.G.: was involved in the design of the study, analysis of data, conception and writing of the manuscript.

## Supplementary Material

Table S1

## Data Availability

RNA-seq data have been deposited at GEO as GSE350898 and are publicly available as of the date of publication. All data reported in this paper may be shared by the corresponding authors upon request. Any additional information required to reanalyze the data reported in this paper is available from the corresponding authors upon request.

## Tables

No main text tables are submitted.

---

**Table S1**

Uploaded as supplemental file.

