## Supplementary figures and images for "TCF21 loss identifies a conserved inflammation-associated fibroblast state across Crohn’s disease and ulcerative colitis"

### Supplemental Figure 1

#### Supplemental Figure 1

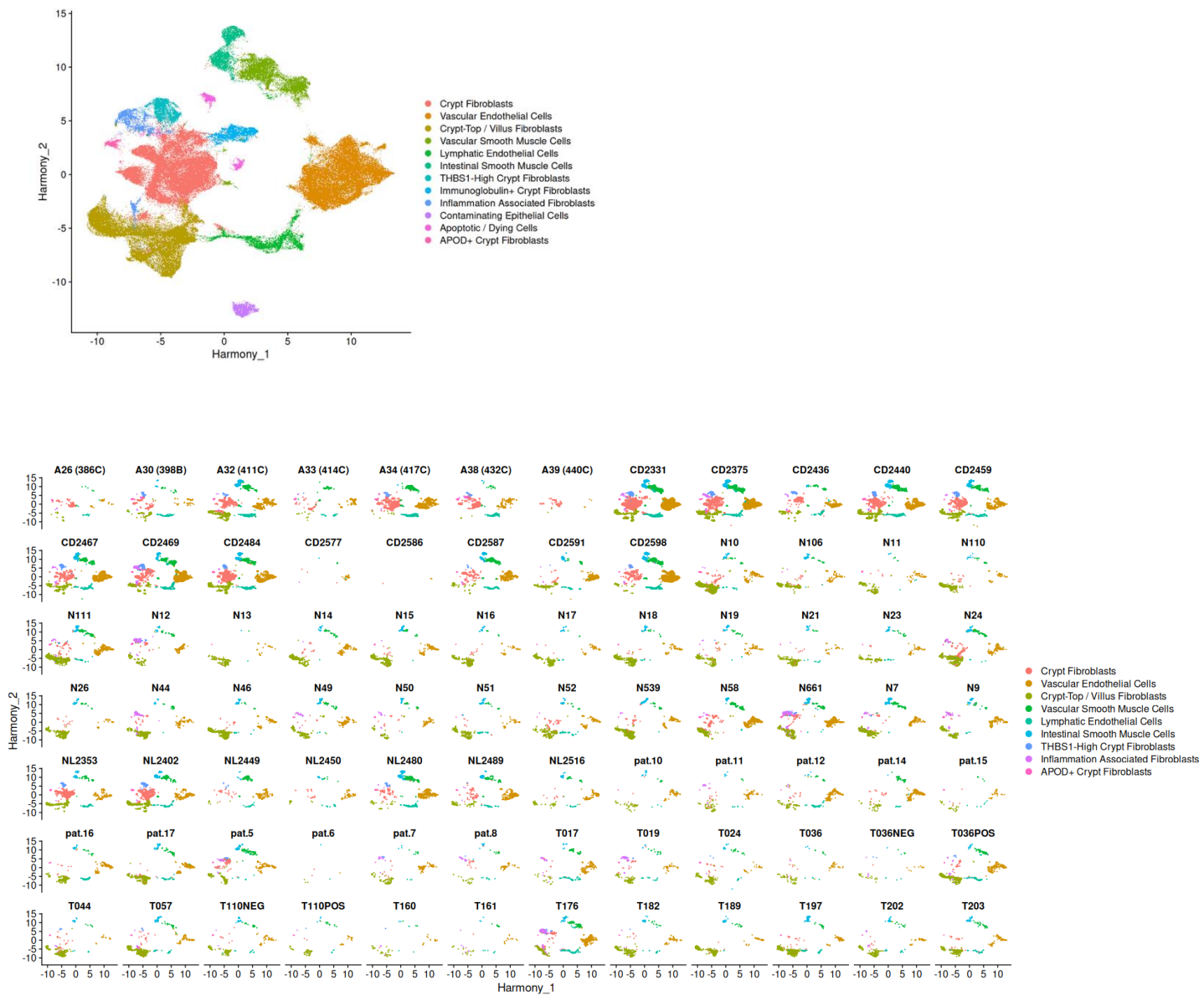
